# Cryptography for genetic material

**DOI:** 10.1101/157685

**Authors:** Sterling Sawaya

**Author notes:** geneinfosec.com.

## Abstract

Genetic information can be highly sensitive and can be used to identify its source. To conceal genetic information, cryptographic methods can be applied to genetic material itself, concealing sensitive information prior to the generation of sequence data. The cryptographic method described here uses randomly divided subsets of barcodes and random pooling to securely generate pools of genetic material. The privacy obtained by these methods are measured here using differential privacy.

## 1 Introduction

Genetic data can contain sensitive health information, such as risk of cancer [Gen94] or neurodegenerative disease [DHMB^+^11, RMW^+^11]. Genetic data can also be used to identify its source (see [HSR^+^08, Cla10, JYW^+^09, VH09, EN14]). Furthermore, when genetic data is combined with other information about an individual, identification of that individual becomes even easier [EN14, HHH^+^15]. This leads to important ethical considerations for those collecting genetic data for research or diagnostics [KBDV^+^10, EWG^+^14].

Consequently, many researchers have turned their attention to protecting genetic privacy. Informed consent can help the participants of a genetic study understand their risks, and frameworks are being developed to help guide scientists who collect genetic information [LCVC08, EWG^+^14]. To allow genetic data to be shared between researchers, various methods of encryption can help prevent re-identification [CMM13, CCL^+^, DDC14a, DDC14b, BBDC^+^11, JWB^+^17, KBLV13, KJLM08, KL15, LGDM10, Mal04, TJW^+^16, XKB^+^14, SSB16].

Those methods are designed for genetic data. The methods described here take a different approach. Here cryptographic methods are applied to genetic material itself, securing genetic information at the molecular level. This approach adds an additional layer of security, allowing genetic material to be sent to untrusted parties for analysis without revealing sensitive information to those parties.

The cryptographic method proposed here utilizes random molecular barcodes, sometimes referred to as unique molecular identifiers or tags. The barcoding of genetic material uses nucleotides as codes. The DNA nucleotides adenine, cytosine, thymine and guanine can be combined in a polymer to form a code. With only four nucleotide bases a large number of codes can be generated. For example, with a nucleotide composed of *n* or fewer bases, 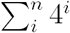 possible codes can be generated. Trillions of possible codes can be generated with short barcodes of only *n ≤* 20.

Various biotechnologies have found applications for random barcodes [SNP^+^16, ZLZ^+^14, SJSX12, SKJ^+^16, LLC^+^16, GBE^+^15, IZJ^+^14, KVK^+^12, BRL^+^15]. These technologies utilize barcoding at the molecular level to improve genetic sequencing accuracy, or examine unique and rare mutations present within a sample. The use of molecular barcoding has yet to be applied to genetic information security. Here, a cryptographic method using random molecular barcodes on genetic material is described.

## 2 Methods

### 2.1 Overview

To begin this crypotgraphic method, genetic material is digested to form separate, unlinked molecules. Second, these molecules are each given random barcodes. Third, the barcoded molecules are combined with other genetic material that has been barcoded, which serve as decoys. Genetic information is concealed because the connections between variants are disrupted, hiding diploid genotypes and haplotypes. Also, the use of decoys can obfuscate which variants in the pool belong to the sample. This general method can be applied in a variety of ways, providing the ability to conceal different forms of genetic information with varying levels of security.

Both randomness and secrecy are required when generating the barcodes. This method takes a set of barcodes, and with secrecy, it randomly divides this set into subsets (Figure 1). Each subset is labelled, and a table of barcodes and labels is provided to the consumer (e.g. Table 1). The consumer can then use the subsets of barcodes to label their genetic samples so that only they know which barcodes belong to which sample.

**Table 1:**
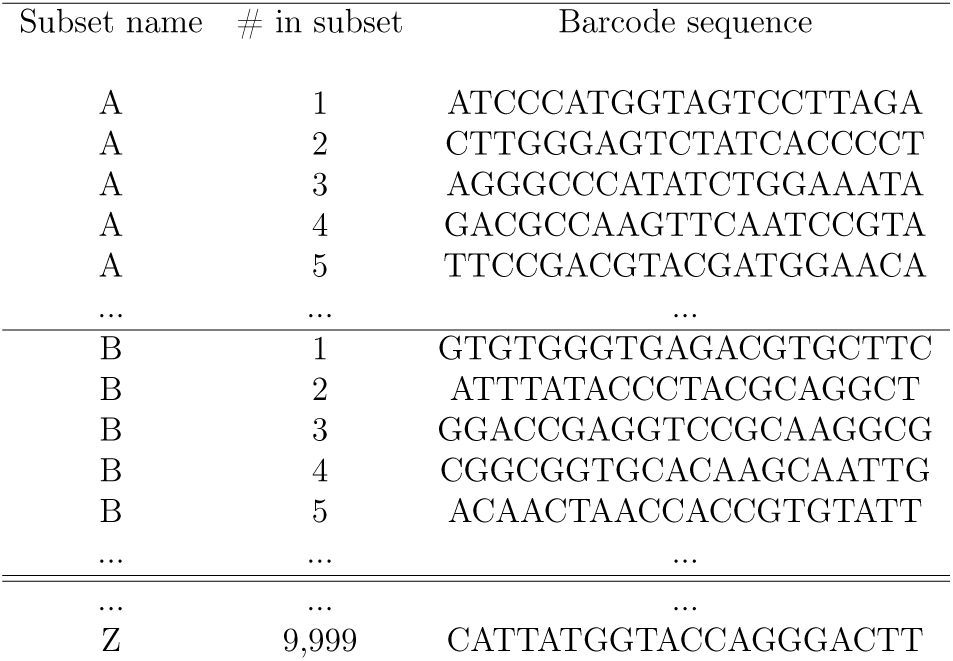
Example of randomized subsets of barcodes. Barcodes are randomly divided into numerous subsets for each consumer. Only the consumer is provided the unique table that can be used to determine which barcodes are in which subsets.

**Figure 1:**
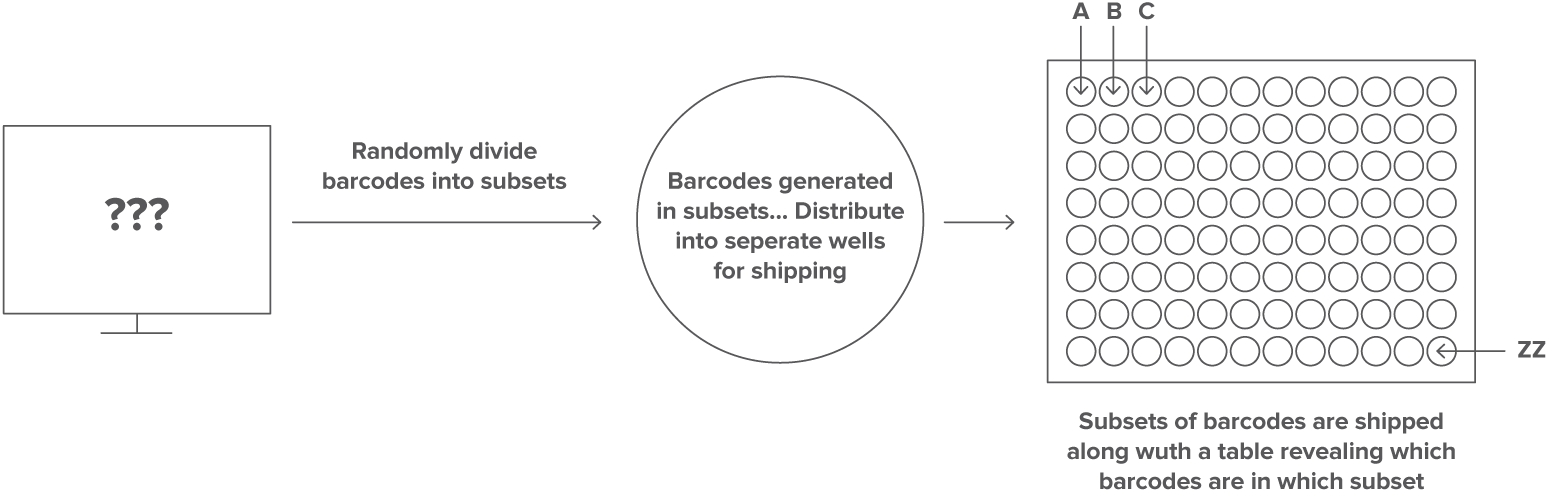
A set of barcodes is randomly divided into subsets. Barcodes are generated in combination with nucleotides used to affix the barcodes to nucleic acids (e.g. adapters). Each subset of barcodes is sealed within a well on a plate. The barcodes present in each subset are only known to the consumer. Barcodes are randomly divided into unique subsets each time they are manufactured.

The barcoding can occur in many ways. Barcodes can be added before or during an enrichment step. Barcoding after or without enrichment can result in a unique barcode for each molecule (Figure 2). If polymerase-chain-reaction (PCR) amplifies the genetic material after it has been barcoded, the resulting molecules would share identical barcodes, indicating they originated from the same molecule and belong to a particular sample (Figure 3). Either way, the barcodes act as random identifiers of the sample. After pooling and analysis, the table of barcodes is required to determine which results belong to which sample.

**Figure 2:**
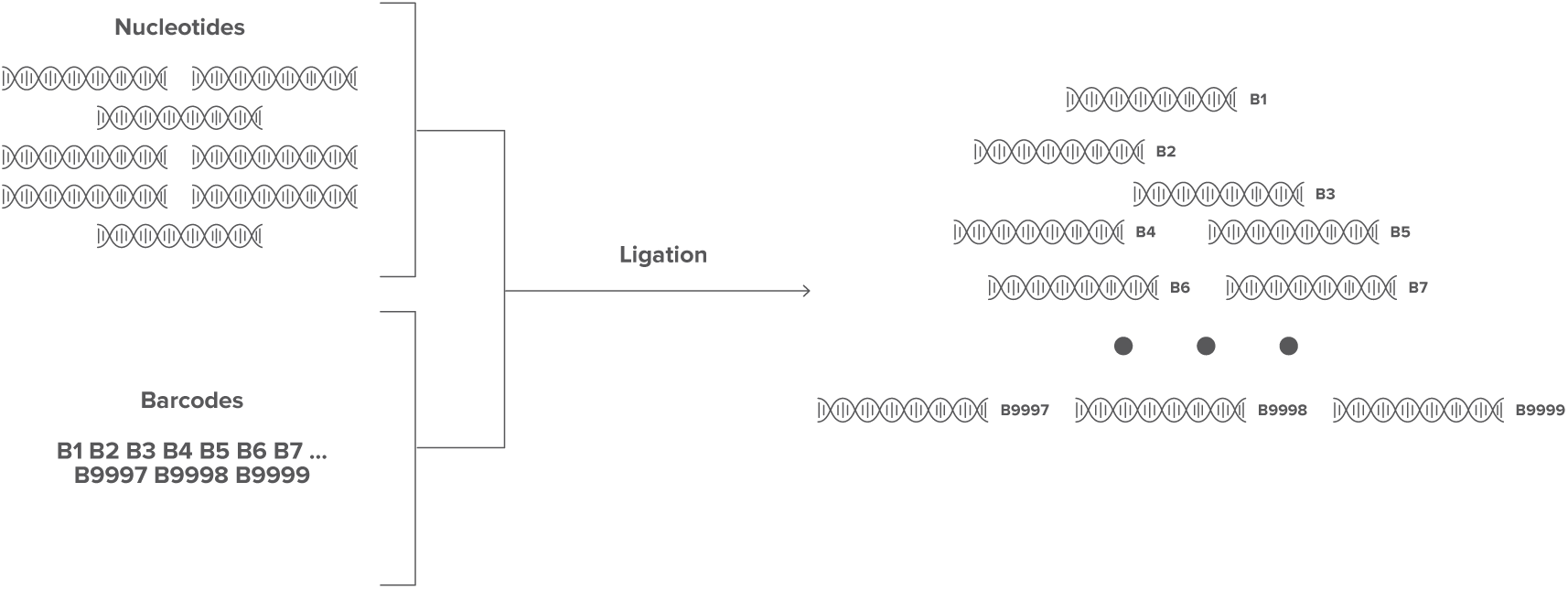
Ligation of barcodes to nucleic acids. A subset Goenfet barcodes is ligated to target nucleic acids, resulting in a unique identifier for each molecule. Only the consumer has knowledge about the barcodes used for a specific sample.

**Figure 3:**
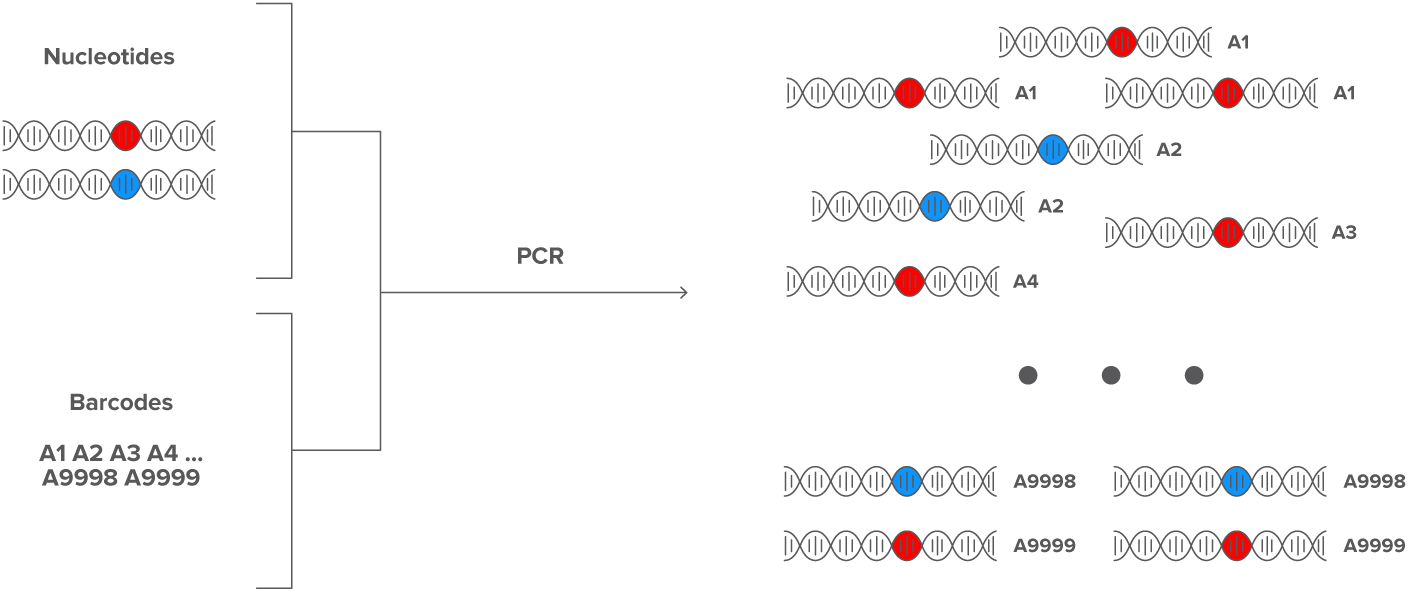
Barcoding before or during enrichment. Nucleic acids are amplified along with barcodes, resulting in multiple nucleic acids with identical barcodes. Nucleic acids that share the same barcode will have originated from the same molecule.

The method of pooling, in which decoys are combined with the sample(s), can be simple or complex, depending on the extent to which one choses to conceal their sample. A simple method of pooling is to barcode a group of samples and combine them together in a pool. These different samples would then act as decoys for each other. More advanced pooling methods can combine samples with non-sample decoys, chosen to conceal specific types of genetic information. The privacy that can be obtained with different pooling methods will be examined here.

### 2.2 Measuring Privacy

The privacy of this process can be examined using the differential privacy method of [Dwo06] using the equation:

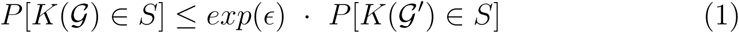

in which ϵ represents the privacy obtained by removing a single individual’s genetic material from 𝒢 to obtain 𝒢 ′ *S* is the range of possible outputs from the process *K*. A group of individuals with genotypes 𝒢 is examined. The genetic material from 𝒢 is converted into genetic data, *K*(𝒢), through process *K*. The probabilities are taken over “coin flips” of *K*.

This randomization procedure differs from those typically used in differential privacy. Here, one can rely on the randomness of a molecular process to obtain privacy, as well as a computer or coin to direct the randomization procedures. This process can be as simple as labeling a group of samples with random barcodes and pooling them together to be sequenced, or more complicated, mutating and amplifying sample genetic material to conceal specific mutations.

To demonstrate how one can measure privacy using this method, consider a scenario in which a small part of the individuals genomes are examined. Assume that only two variants are present within the population in this region. Denote alleles or haplotypes as “*A*” for the common variant and “*a*” for the alternative, less frequent variant. The frequency of the less frequent variant, denoted as *p*, is thus on (0, 0.5]. The frequency within the genetic data, however, is on [0, 1], depending on the genotypes of the individuals being analyzed as well as the process by which the genetic data is generated. Here, the genotypes of the individuals in the pool are assumed to be in Hardy-Weinberg equilibrium.

Differential privacy methods are applied to a wide variety of data-sets, and variety of methods can be used to estimate differential privacy on these different data (e.g. [WLF16, BNS^+^16, BD14]). Many methods examine the sensitivity of the randomization procedure, the maximum difference in output that can be generated [Dwo08], as:

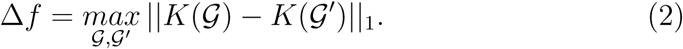

Denote 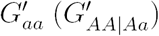 as the pool of genetic material with the genetic material of an individual’s with genotype *aa* (*AA* or *Aa*) removed. For many of the procedures examined here, removal of homozygous rare alleles results in the largest change in the output. That is, it can be shown that

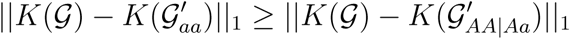

for many procedures *K*. Consequently, concealing genetic information for individuals with uncommon genotypes is more difficult than concealing that of those with more common genotypes. Therefore, measuring the privacy obtained by removing an individual with the *aa* genotype determines the value of ϵ. If the frequency of the rare allele is small enough that no *aa* individuals are present in the pool, privacy is measured with 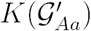.

With the above considerations, equation [1] can be restated as:

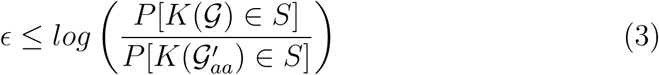

for many procedures *K*. This can lead to a comparison between KullbackLeibler Divergence (KLD) [Kul97] and differential privacy [WLF16, BNS^+^16, BD14]. The KLD between two distributions is a measure of the information gained if one distribution is used in place of the other. For two distributions, P and Q:

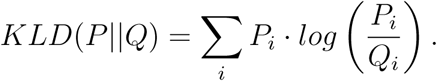

This can be related to privacy measured in [3], as the *E*(ϵ) is the same as 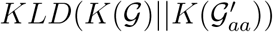, which has been termed “On-Average KL-Privacy” [WLF16]. Therefore, the differential privacy measured here can be interpreted as the difference in information between the entire pool of genetic material, and the pool with the *aa* genotype individual’s genetic material removed (from the perspective of the original pool). That is, the information lost if *K*(𝒢) is encoded by 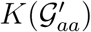.

Consider a pool of genetic material from *N* individuals that has been amplified *X* times, resulting in 2*NX* alleles in the pool. Simply pooling together genetic material and sequencing a portion of that pool is a randomization procedure that offers privacy. Sampling from a well mixed pool results in data for each allele that follows a hypergeometric distribution. If a sequencing method provides *y* total sequences, then the probability of *z* sequences with the “*a*” variant, *P*_*z*_(*a*), from a pool of size 2*NX* is:

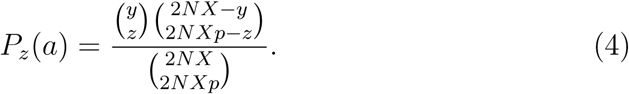

To measure privacy in this method, compare (4) with the probability of sequence results from the same pool with one individual with *aa* genotype removed, resulting in:

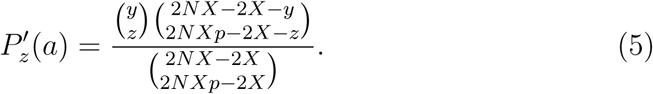

Measuring the KLD between these two distributions provides the expected KL-privacy for *aa* genotypes, setting the bound ϵ:

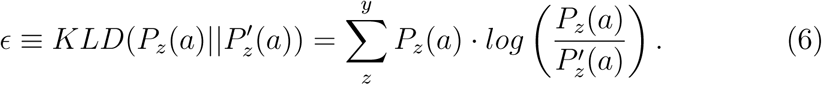

The variables here have significance in a genetic sequencing analysis, and thus their values must be chosen carefully so that the proposed analysis would be reasonable. The number of times an allele from an individual is sequenced in a pool, *P*_*z*_(*allele*), is also a hypergeometric function determined by the values of *X* and *y* chosen for a pool of *N* individuals,

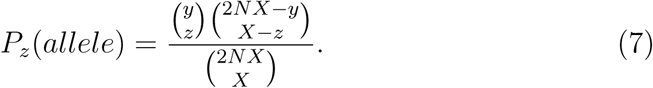

Sequencing analyses are designed so that each allele from every individual achieves an appropriate number of reads. The average number of times an individual’s allele will be sequenced is *y/*2*N*. This expectation, however, may not be likely in many designs, and a more appropriate design would use (7) to ensure that each individual receives sufficient coverage with an appropriate probability.

### 2.3 Advanced methods

First, consider that the amplification amount, *X*, can be a random variable. The randomness can be directed by a computer or by flips of a coin, randomly selecting an amount for each sample, *X*_*i*_, to be added to the pool. Now, consider that the amplification occurs by PCR, and this process is intrinsically random. The randomness of amplification can be further randomized by a computer. For example, a computer can provide a random number of cycles of PCR by which each sample is amplified, or a random quantity of the various PCR ingredients to further vary the amount by which each sample is amplified.

Any random amplification procedure results in the *X* becoming a random variable. Here, the random amplification is applied to each individual sample, such that each allele in the sample receives the same amplification. Consequently, the privacy measured in (6) must then be measured over the possible values of *X*. The addition of randomness to the process in the amplification can increase privacy provided by this method. In fact, due to the imprecision of aliquoting genetic material, as well as the randomness that occurs in PCR, one may consider *X* to always be random. To help estimate the privacy the randomness of PCR can be modeled, e.g. [JK03, Pia04, YY09, LJJ05].

Mutation can also be utilized for privacy. Mutations randomly occur during PCR amplification [KZ15, PO17], with some polymerases having higher mutation rates, for various different types of mutation [PO17]. Furthermore, site directed mutagenesis, e.g. [Car86, HAT^+^89, HHH^+^89, LGH90, WCS^+^94, KM97], can be utilized to obtain specific mutations from the sample. That is, a proportion of the sample can be (randomly) mutated to become decoys with specific variations to be added to the pool. Importantly, the mutated genetic material must be labelled uniquely and combined with uniquely labelled, non-mutated material, so that one can determine which sequencing results belong to the non-mutated genetic material. Methods using mutation can add additional sequencing costs because the sequencing of mutated genetic material usually does not provide useful information to the consumer. However, as sequencing costs continue to decrease, processes that include mutation will become increasingly cost effective.

Applying a random mutation step alters the equations used to estimate privacy. The total size of the pool of genetic material can be generalized, represented here by the variable *Z*. The pool *Z* can then be divided into the quantities of the separate alleles, here 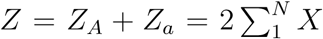. The number of specific alleles in the pool is the sum of the contribution from each genotype. For *Z*_*a*_:

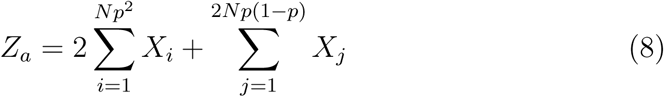

and similarly for *Z*_*A*_:

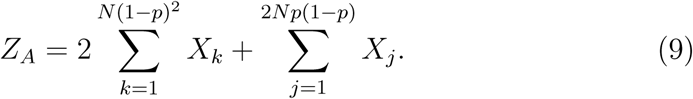

The hypergeometric (4) then becomes:

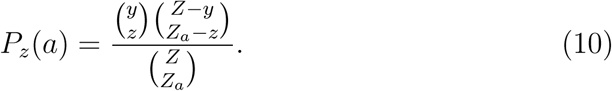

Denote 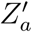 as the number of “*a*” alleles in a pool from which an individual with genotype *aa* has been removed:

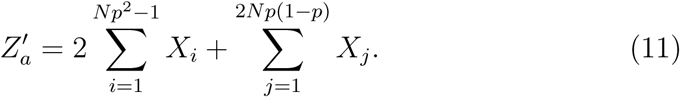

The total resulting pool size for this pool, *Z′*, is the sum 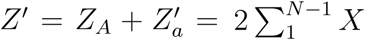. With the new variables for the size of the altered pool and quantity of “*a*” alleles in the pool, (5) then can be generalized as:

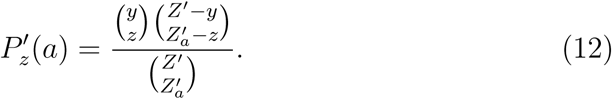

If mutations are considered, then the final pool *Z* can be modeled as a mixture of alleles that have been replicated and mutated from an original pool of samples. Denote the mutations between alleles as:

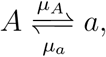

If the mutation between the two variants are equal (*µ*_*a*_ = *µ*_*A*_), then the number of “*a*” alleles in the pool is:

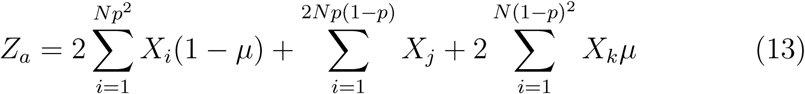

and the number of “*A*” alleles is:

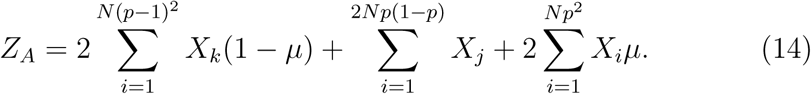

Now consider the pool in which an individual with genotype *aa* has had their genetic material removed. Denote this comparison pool as 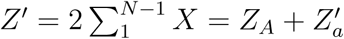 and its quantity of “*a*” alleles as 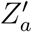, then:

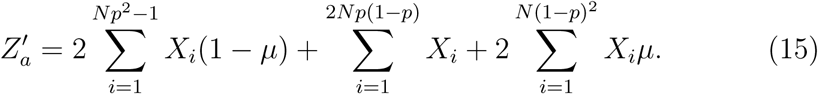

As before, privacy is measured by comparing (10) and (12) using (6).

## 3 Results

### 3.1 Non-random pooling

Privacy is measured for pools of varying numbers of individual samples, each sample pooled with equal proportions (Figure 5). Populations, *N*, of 400, 800 and 1,600 individuals, among a range of allele frequencies are examined, with the amplification *X* = 10^6^. The number of reads is 8,000, 16,000 and 64,000 for 400, 800 and 1,600 individuals, respectively, providing a *P*_*z≥*5_(*allele*) *>* 0.97, with an average of 10 reads per allele.

**Figure 4:**
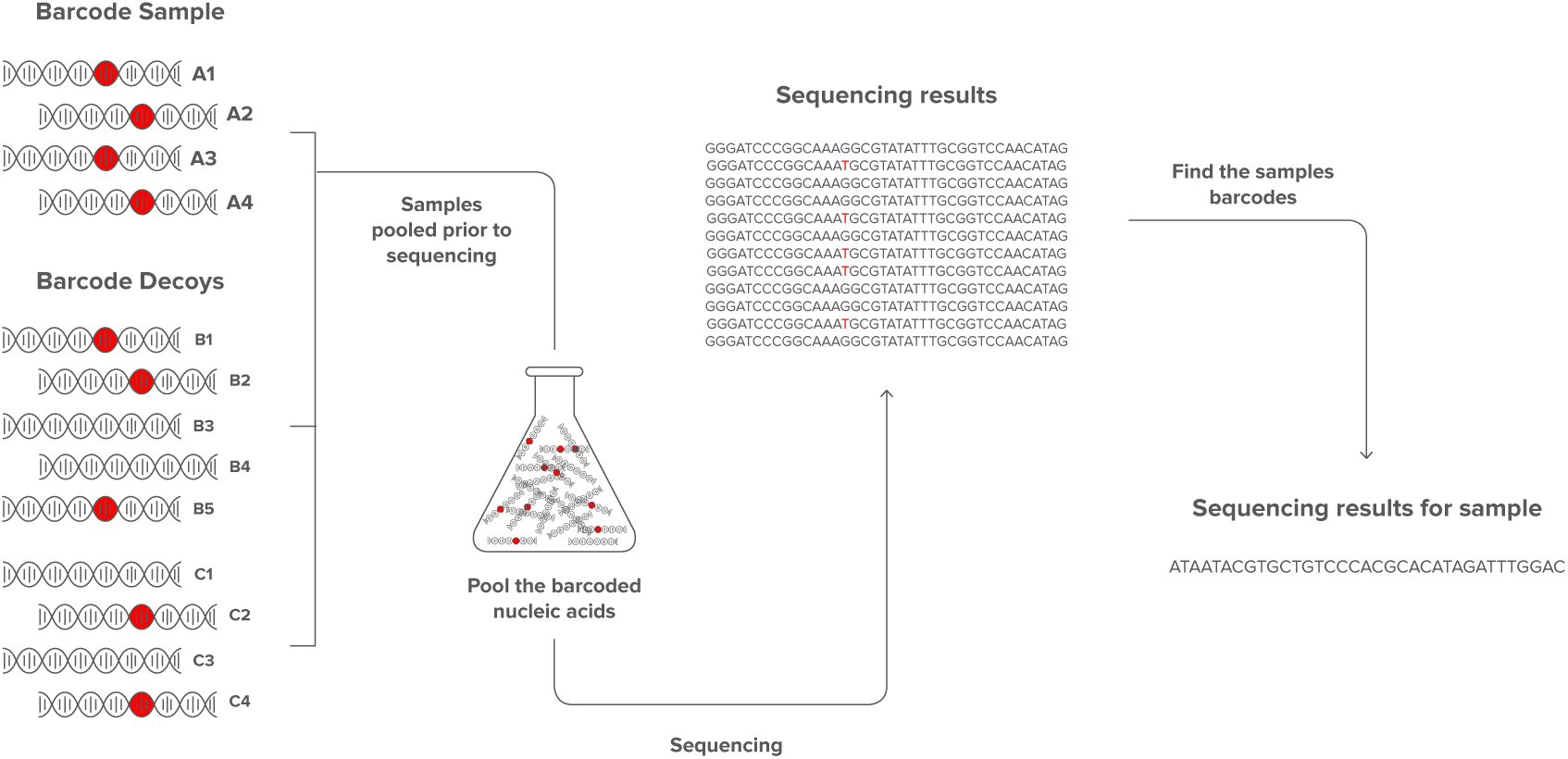
Pooling barcoded nucleic acids. The final step to concealing genetic information is the pooling of sample nucleic acids with decoy nucleic acids. A table of barcodes can be used to determine which sequences belong to which samples.

**Figure 5:**
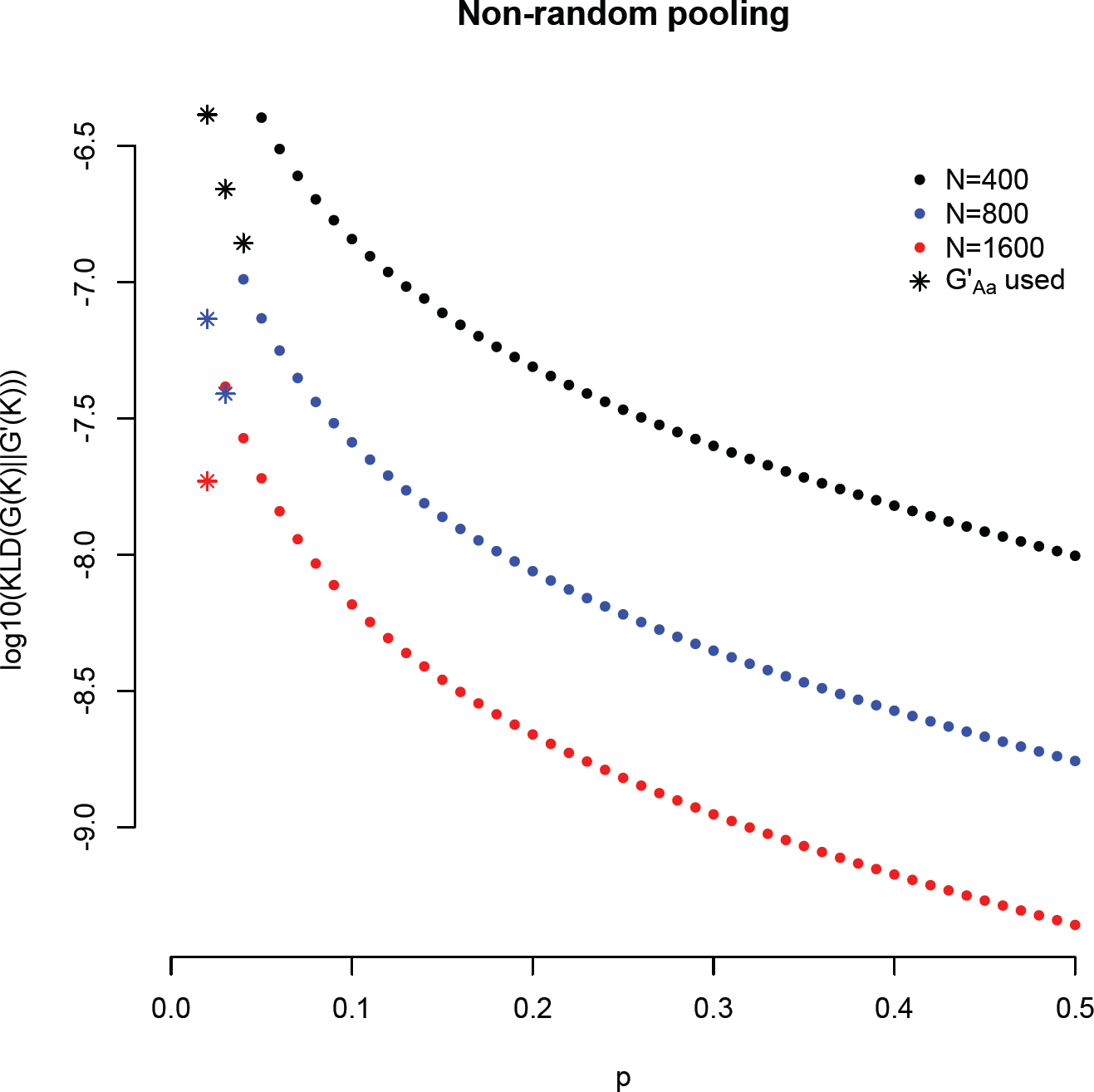
Privacy obtained by pooling samples together. The log (base 10) values of *E* for different pool sizes and different allele frequencies. Samples are pooled together in equal proportions for N=400 (black), N=800 (blue), and N=1600 (red). Samples were amplified to have 10^6^ molecules prior to pooling, and the pool was sequenced 20. *N* such that the average number of sequences per allele is 10. If the allele frequency is small enough that zero “*aa*” genotypes are expected in the pool, then privacy is measured by removing an “*Aa*” genotype individual from the pool instead of aa individuals (points plotted with *\**).

### 3.2 Random pooling with mutation

To examine random pooling and mutation, a population of 100 samples is used (Figure 6). For comparison, non-random pooling of each 100 samples is measured (black points). For random pooling, each individual has 1,000 of each of their alleles added to the pool, and then, for each sample, a coin is flipped two times, and an additional 1,000 of each allele is added for every flip that landed heads (red points). The resulting quantity of each allele for each sample in the pool follows a binomial distribution. The same random amplification method then is applied, but with 20% (orange points) and 40% (blue points) of the alleles mutated to the other variant. Random pooling provides more privacy than non-random pooling, and privacy is further increased if a mutation step is applied.

**Figure 6:**
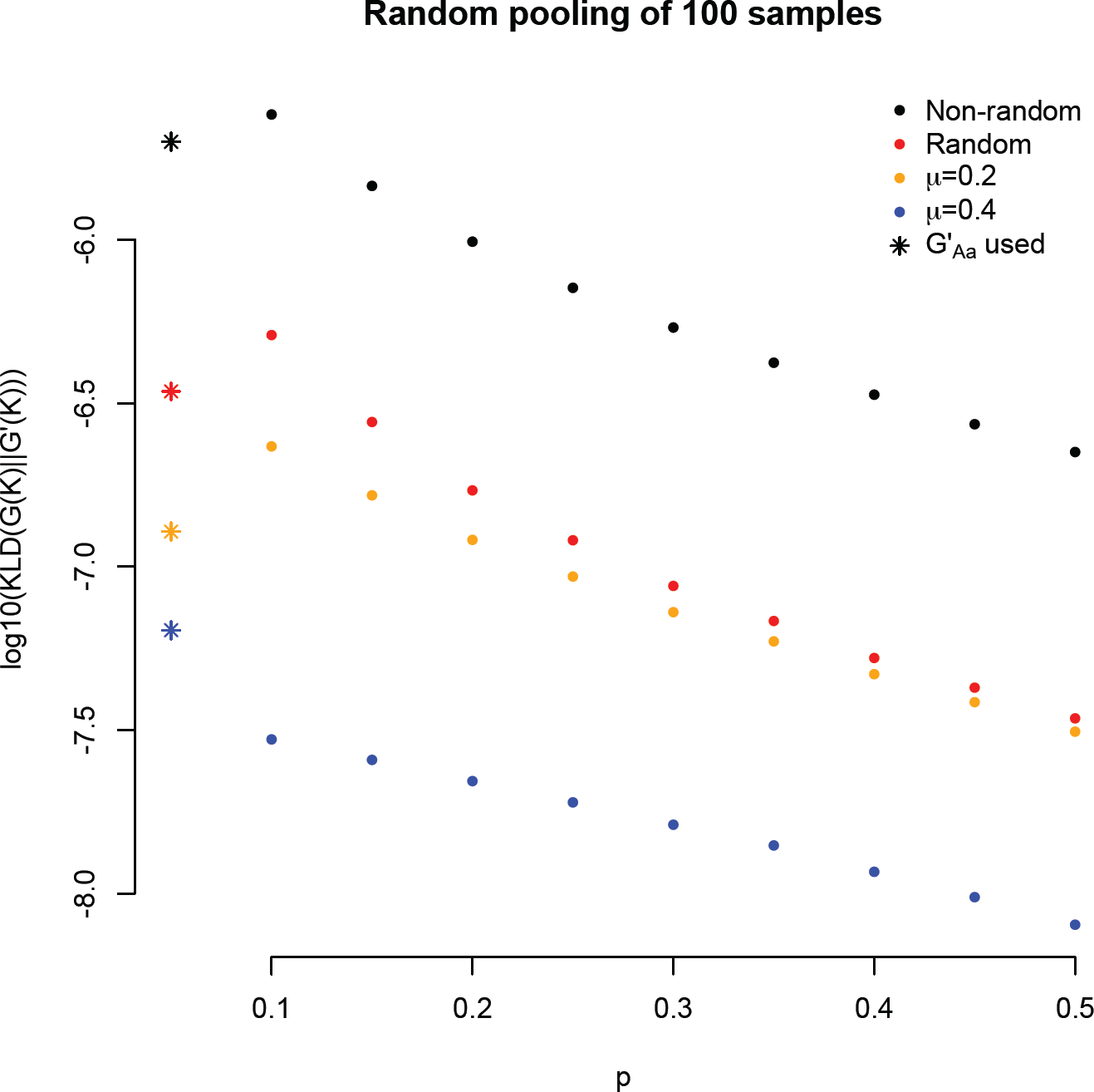
Privacy obtained by randomly pooling samples together with mutation. The log (base 10) values of ϵ for different pool sizes, different allele frequencies, for different pooling methods. Black points represent pooling samples of 100 individuals in equal, non-random proportions. Red points represent a random pooling of 100 individuals (see text for details). Orange (blue) points represent a random pooling of 100 individuals with a mutation rate, *µ*, of 0.2 (0.4). If the allele frequency is small enough that zero “*aa*” genotypes are expected in the pool, then privacy is measured by removing an “*Aa*” genotype individual from the pool instead of aa individuals (points plotted with *\**).

A random selection of 4,000 reads is then obtained from the pool, resulting in an average of 20 reads per individual when a mutation step is not applied. With a mutation step, some individuals receive far fewer reads and some receive far more (Figure 7). Furthermore, the mutated alleles, which have been uniquely tagged to indicate they are mutants, do not necessarily provide useful information about the sample. Consequently, pooling procedures which apply mutations result in fewer informative reads.

**Figure 7:**
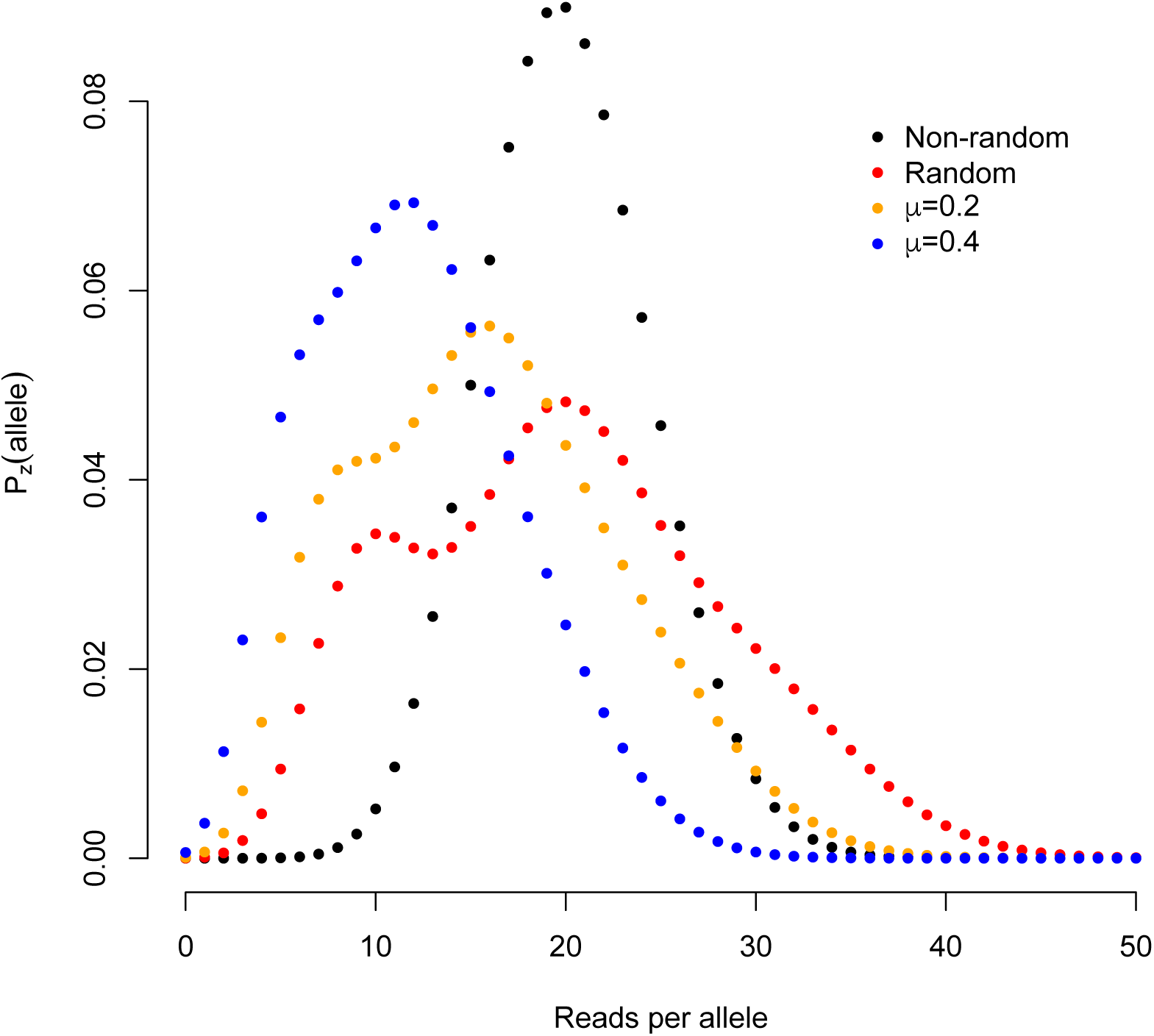
Probability of read number for sample alleles for different pooling methods. The number of reads for non-random pooling (black points) approximately follows a binomial distribution. Red points represent a random pooling of 100 individuals. Orange and blue points are random amplification, with a mutation step applied. The reads of mutated alleles are not counted, as they are not typically informative.

## 4 Discussion

The statistics presented here demonstrate that genetic information can be concealed with the methods proposed. The proposed methods have a wide range of potential applications in genetics, and are not limited to those examined in the results section. For example, the methods can be applied to more than one genetic loci, or to genetic loci that that have more than two states, such as tandem repeats. The use of differential privacy to measure the expected privacy of these methods can also be applied to these other applications. Alternatively, detailed models of adversarial knowledge can also be used to estimate privacy.

Any application of these methods requires careful attention to how information exists within target genetic material and how it can be concealed. Because this method does not necessarily conceal all information present within genetic material, the consumer needs to decide which information they conceal and how they conceal it. For example, the basic application of these methods does not conceal the allele frequency in the samples. Advanced applications can conceal the allele frequencies, but may require more sequencing reads to obtain the same amount of information. Importantly, due to correlations between genetic variants, applying these methods to multiple loci requires special considerations to estimate differential privacy (see [CYXX16, KM11] for estimating differential privacy with correlated data).

The advanced methods presented in the results section can be applied with readily available technology, a coin to randomly determine the quantity of genetic material to add to the pool and a controlled quantity of mutated alleles obtained through site directed mutagenesis. However, one can consider all molecular genetics lab work to have a degree of randomness (even when randomness is not desired). Consequently, all applications of this method will have more randomness, and thus usually more privacy, than the ideal applications measured here. Modeling PCR randomness to estimate privacy requires a detailed understanding of genetic information, DNA/RNA replication, as well as mutation rates. However, for many applications of this technology non-random pooling may be sufficient, allowing for relatively straightforward measures of privacy.

Ultimately, the details of the cryptographic method used by any consumer of this technology can be kept secret, and can be altered when applied to different groups of samples. This allows the consumer to control how they conceal their genetic information, further inhibiting potential adversaries from extracting useful data from the sequencing results. The appropriate use of securely generated random barcodes allows sensitive genetic information to be concealed within genetic material, securing it at its source. With hope, these methods will permit the collection of genetic information without jeopardizing sensitive information.

## Acknowledgement

Thanks to Dr. Dylan Albrecht for helpful discussions on some of the statistics.

